# Cooperative hydrodynamics accompany multicellular-like colonial organization in the unicellular ciliate *Stentor*

**DOI:** 10.1101/2023.01.10.523506

**Authors:** Shashank Shekhar, Hanliang Guo, Sean P. Colin, Wallace Marshall, Eva Kanso, John H. Costello

## Abstract

Evolution of multicellularity from early unicellular ancestors is arguably one of the most important transitions since the origin of life^1,2^. Multicellularity is often associated with higher nutrient uptake^3^, better defense against predation, cell specialization and better division of labor^4^. While many single-celled organisms exhibit both solitary and colonial existence^3,5,6^, the organizing principles governing the transition and the benefits endowed are less clear. Using the suspension-feeding unicellular protist *Stentor coeruleus*, we show that hydrodynamic coupling between proximal neighbors results in faster feeding flows that depend on the separation between individuals. Moreover, we find that the accrued benefits in feeding current enhancement are typically asymmetric– individuals with slower solitary currents gain more from partnering than those with faster currents. We find that colony-formation is ephemeral in *Stentor* and individuals in colonies are highly dynamic unlike other colony-forming organisms like *Volvox carteri* ^3^. Our results demonstrate benefits endowed by the colonial organization in a simple unicellular organism and can potentially provide fundamental insights into the selective forces favoring early evolution of multicellular organization.

---

Suspension-feeding unicellular protists inhabit a fluid world dominated by viscous forces that limit prey transport for feeding ^3,7,8^. Using either flagella or cilia, many of these organisms generate microcurrents that actively transport dissolved nutrients and smaller prey critical for their nutrition ^3,9,10^. A protist’s ability to favorably alter its feeding current so as to enhance feeding rate would therefore be beneficial to its survival. Can colony formation enable unicellular protists to enhance their feeding flows? Colonial protists have been suggested to generate stronger flows by combining individual feeding microcurrents of neighboring colony members. Colony forming protists can broadly be classified into two categories depending on presence (or absence) of physical linkages between colony members. A relatively smaller number of organisms, including *Volvox carteri* ^3^ and the choanoflagellate *Salpingoeca rosetta* ^6^, form colonies where members are physically attached to each other. Volvox forms spherical colonies with individual cells embedded within an extracellular matrix jointly secreted by identical, clonal colony members. The combined flows can act over longer distances ^11^ and have been proposed to transport greater amounts of fluid per individual protist ^11,12^. This has been suggested to enable Volvox colony members to grow and reproduce faster^13,14^.

However, if we step backwards, even before these more advanced examples, the majority of known colony-forming protists form colonies by aggregating in relatively high numbers with no observable linkages between colony members. This more basal condition mimicking multicellular-like behavior precedes the evolution of an organizing extracellular matrix or cell-cell junctions, as seen in volvox or choanoflagellates^4^. We therefore asked if individuals in such a loosely organized colony are capable of coordinating their activity to achieve common goals, or is such cooperation an exclusive province of colonies in which individuals are physically connected?

To answer this question, we specifically address two issues: the positioning of individuals in a colony with no common extracellular matrix and the hydrodynamic consequences of such colony formation. We chose the ciliated unicellular protist *Stentor coeruleus* to examine these questions. *Stentor* individuals generate feeding currents by beating band of cilia near their “heads” at their anterior end and attach on organic surfaces in the wild via an organelle called “holdfast” at their posterior end^15^ (Fig. 1a-b). *S. coeruleus* occur as freely swimming individuals under low prey conditions. In higher food conditions however, *Stentor* individuals can reversibly aggregate into hemispherical colonies by anchoring themselves on a surface in close proximity of each other. Colony members can sway their feeding apparatus (i.e., heads) back and forth without detaching their holdfast from surfaces (see Movie 1). We examine the hydrodynamic cooperation between these unconnected colony members and find that although benefits accrue for neighboring individuals, the benefits are typically asymmetric between individuals and that hydrodynamic cooperation is highly promiscuous among colony members.

**Fig. 1:**
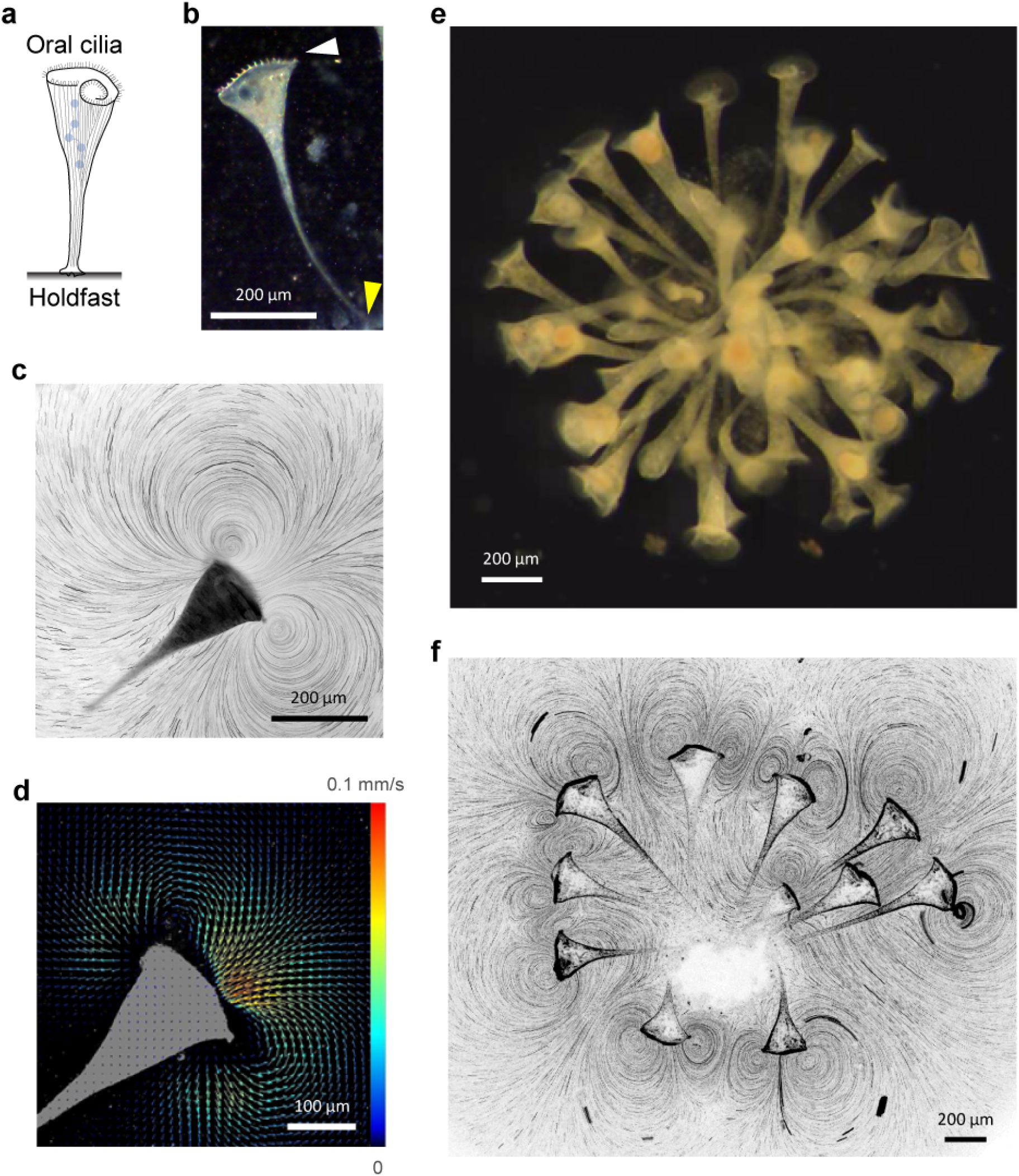
Ciliary flowfields generated by solitary and colonial *Stentor species*. **(a)** Schematic representation of a *Stentor coeruleus* attached to a surface. The oral ciliary band is located at the anterior (head) and forms the feeding apparatus. The attachment organelle “holdfast” is located at the posterior of the organism (adapted from ^16^). **(b)** Darkfield image of a *S. coeruleus* individual anchored on the glass coverslip. The white and yellow arrowheads indicate the ciliary band and the holdfast respectively. **(c)** Tracer particle tracks from the time-lapse recording of the flow generated by a *S. coeruleus* individual. The dark shape shows the location of the organism. **(d)** Particle image velocimetry analysis of the flow generated the organism shown in (c). The direction of the arrows denotes the local flow direction and color denotes the magnitude of the flow speed. **(e)** Self-assembled colony formed by wild *S. muelleri* suspended in solution (see Movie 1) **(f)** Tracer particle tracks from the time-lapse recording of the flow generated by a colony of 11 *S. coeruleus* individuals. Outlines of individual colony members can be seen.

Individual *Stentor coeruleus* freely suspended in water were incubated either alone or in larger groups in a PDMS chamber. Once anchored on the PDMS-attached glass coverslip by their holdfast, *S. coeruleus* generate feeding currents by beating their oral cilia in a metachronal wave (Fig. 1c) ^16,17^. Dilute amounts of whole milk or microparticles added as flow tracer enabled direct visualization of flow streamlines under dark field microscopy ^18,19^. Tracer trajectories confirmed laminar flow consistent with the estimated low Reynolds number (∼0.1) for an organism about the size of *S. coeruleus* (∼1 mm) swimming in water. The flowfield extended well beyond the organism’s size and was characterized by appearance of two symmetric regions of vorticity (these appear as vortices when observed in 2D, and are therefore referred to as “vortices” henceforth) with opposing spin on either side of the oral opening (Fig. 1c). Since *S. coeruleus* are found attached to organic matter in their natural habitat (i.e. aquatic bodies like ponds), their attachment on glass coverslip essentially recapitulates a no-slip boundary they attach to in the wild ^20^. Feeding current velocities were determined by micro particle image velocimetry (μPIV) of high speed video recordings of the flows (Fig. 1d) ^21^. Velocities were the highest (∼100 μm/s) near the oral apparatus and the magnitude tapered off by an order of magnitude over a distance equivalent to organismal size (∼1 mm). Since only the orthogonal portion of the flow between the two vortices reaches the oral opening and can be productively filtered for prey, we define the feeding flow velocity (U) as the downward velocity component averaged over the ciliated ring area. Additionally, U was measured 0.25 mm away from the ciliated ring to prevent contamination of the PIV analysis due to its inability to separate tracer movements from ciliary beating (Extended Data Fig. 1). The average feeding current velocity of solitary individuals was found to be about 70 µm/s (n=16) (Extended Data Fig. 2), with at least a 3-fold difference between the strongest and weakest solitary flows.

When multiple individuals were incubated in the same chamber, the organisms readily self-assembled into colonies exhibiting radial symmetry around the point of anchoring. This behavior was conserved in both of the *Stentor* species we examined – *S. muelleri* which were acquired from their native environment (Fig. 1e) and the lab cultured *S. coeruleus* (Fig. 1f). Unlike solitary individuals, *S. coeruleus* colonies exhibited flowfields with large number of complex interfering vortices (Fig. 1f) possibly due to crosstalk between solitary flows of proximal individuals. We therefore investigated if this reduced proximity between individuals in a colony led to enhancement of their feeding flows.

To test this, we employed a simplified *S. coeruleus* pair system. Two individuals were incubated simultaneously in the PDMS chamber. Once attached on the glass coverslip, while the individuals didn’t alter their point of anchoring, their heads (i.e. oral openings) exhibited movement with respect to each other. As a result, the exact separation between the two heads varied over time. The appearance of the combined flowfield also changed as a function of the distance between the two heads (Fig. 2a). As the separation gradually reduced, their otherwise independent flows began interacting and the independent vortical structures of the two organisms became asymmetric i.e. the size of the inner vortical structures were smaller than those of the outer ones (Fig. 2a). As the inter-individual separation was gradually eliminated (i.e. Δ∼0), only two external vortices remained. The flowfield at low separation resembled a flowfield expected from a *S. coeruleus* much larger than either of the two alone. PIV analysis confirmed that changes in flowfield appearance were reflected in changes in flow velocity as well (Fig. 2b). The two flows with velocities 120 and 190 μm/s merge into a single faster flow with a velocity about 330 μm/s as the separation between the two individuals is eliminated (Δ=0) (Fig. 2b,c), translating to an asymmetric benefit of 2.8x and 1.7x enhancement for the flow reaching the ciliated oral discs of the two individuals. Thus, although both the weaker and the stronger organism appear to benefit from their close proximity to each other, the advantage each gains from the pairing is dependent upon their initial flows i.e. flow enhancement of the two individuals is asymmetric. This pattern was observed for multiple different organismal pairs (Fig. 2d). The weaker organism gained more from the pairing than the stronger one. We found no evidence that the frequency of the ciliary waves was altered as separation (Δ) between individuals changed (Extended Data Fig. 3). Whether the increase in rate of feeding current translates into more prey capture has been difficult to test due to *Stentor*’s sensitivity to bright light in the visible spectrum. This has thus far prevented us from carrying out quantitative imaging of prey capture.

**Fig. 2:**
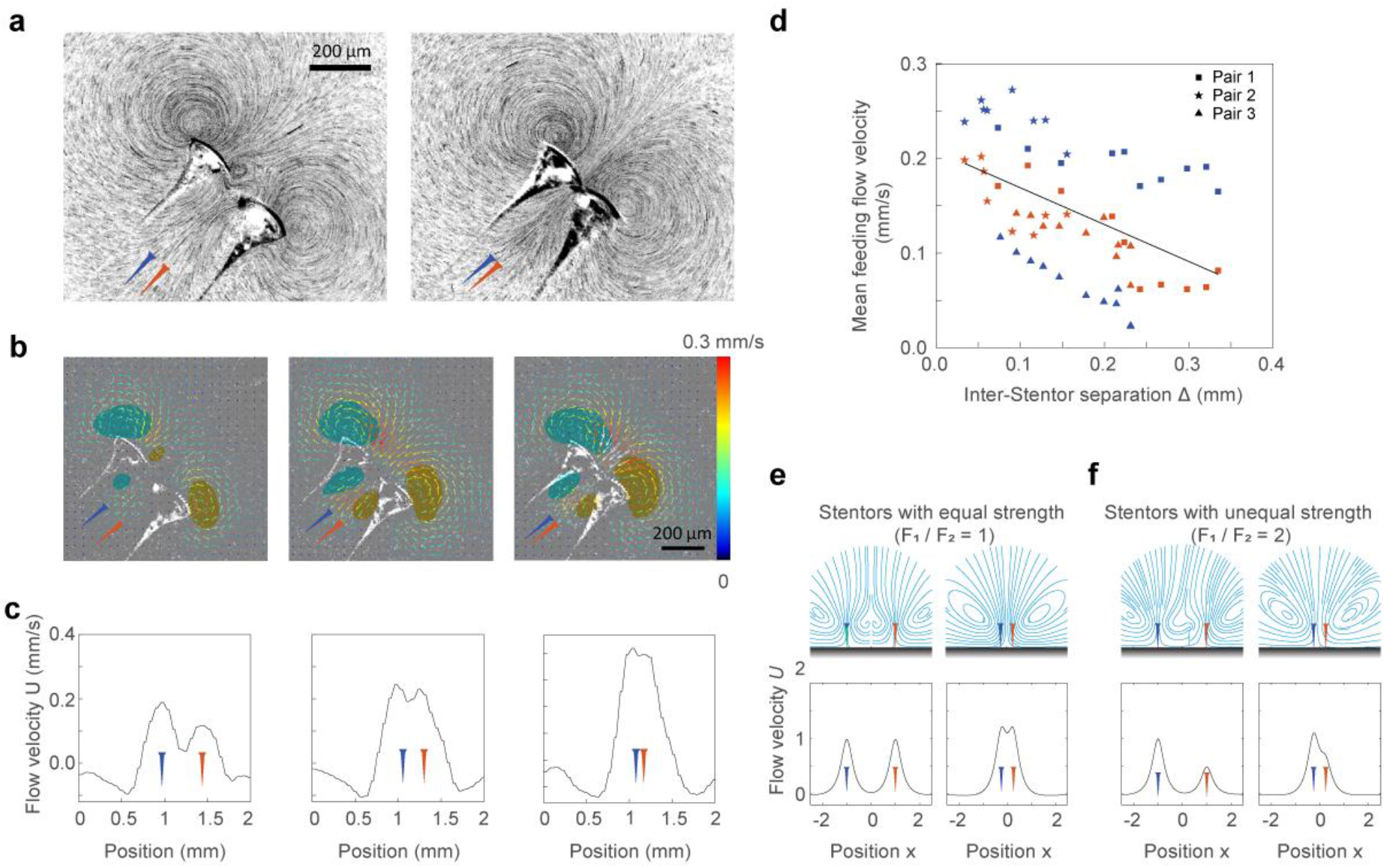
Magnitude of feeding flow of a *S. coeruleus* pair increases with decreasing inter-organismal separation. **(a)** Tracer particle tracks of the flowfield generated by a pair of *S. coeruleus* at varying separation distance (Δ). Left: Intermediate separation. In addition to the two outer vortices, two smaller internal vortices are observed. Right: Zero separation between the organisms. Only two outer vortices are seen. Insets: Schematic representation of the two individuals (red and blue). **(b)** Particle image velocimetry analysis of the combined flow generated by a pair of organisms as a function of decreasing (from left to right) inter-organismal separation. The direction of the arrows denotes the local flow direction and color denotes the magnitude of the flow speed. The cyan and green bubbles proximal to the oral opening represent flow directionality: clockwise (cyan) and counterclockwise (dark green). **(c)** Vertical velocity profile of the combined flowfield of the pair shown in (b) at different separations corresponding to the images shown in (b). Insets: Schematic representation of the location of the two individuals. **(d)** Mean feeding velocity measured for 6 different *S. coeruleus* individuals (across 3 pairs) as a function of the inter-organismal separation. Blue and red symbol color represent left and right organism of each pair. Pair 1 are the organism shown in panels (a-c) Thick black line is a linear fit of the combined data of 6 *S. coeruleus* individuals. **(e)** Fluid flows generated by a *Stentor* pair, modeled as a pair of regularized Stokeslets near a wall with force ratio *F*_*1*_/*F*_*2*_ = 1, inter-stentor separation distance Δ = 2 (left) and Δ = 1 (right) **(f)** Force ratio *F*_*1*_/*F*_*2*_ = 2 and inter-stentor separation distances Δ = 2 (left), and Δ = 1 (right). Top row: flow streamlines. Bottom row: downward velocity component of the flow velocity measured at a vertical distance (*h*/*H*) = 1/4 from the location of the Stokeslet forces.

To test whether the enhanced flow stemmed from hydrodynamic cooperation between individual flows, we modelled the *S. coeruleus* pair as two regularized stokeslets pointing downwards placed side-by-side near a wall (Fig. 2e) ^20,22-25^. In this simple Stokes flow model, the details of the cilia length, beating frequency and waveform of *Stentor* individuals are subsumed into a single averaged parameter - the force *F* that an individual *Stentor* exerts on the fluid. Dimensional analysis shows that the characteristic magnitude of this force scales as *F* ∼ *ηUH*, where *η* is the fluid viscosity, *H* is the length of a typical *Stentor* individual (∼1 mm) and *U* is the typical flow speed (∼100 µm/s). Our model faithfully reproduced the flowfields created by a pair of *Stentor* individuals (Fig. 2e). Interestingly, our model predicted that both the appearance as well as the velocity profile of the combined flowfield would depend upon two main factors: the strengths of the two individual *Stentor*s (*F*_1_ and *F*_2_) and the distance of their separation (Δ) (Fig. 2e). For pairs in which the members have equal strength (i.e., force ratio *F*_1_/*F*_2_ = 1), the predicted flowfield is left-right symmetric for all values of Δ. However, when the two *Stentor* individuals have unequal strengths we see asymmetries in the flowfield.

How do these asymmetries affect feeding flows for interacting *Stentor* individuals? We used our mathematical model to predict the resulting average feeding flows of *Stentor* pairs comprised of organisms with equal and unequal strengths. In both cases, we found the mean flow for the pair increases as the separation distance between the two individuals decreases. For the pair of equal strength (*F*_1_ = *F*_2_), the average feeding flow to each *Stentor* is identical (Fig. 2e, 3a). Interestingly, when the individual *Stentor*s in a pair have unequal strength, the resulting feeding flows to each *Stentor* are not identical (Fig. 3a,b). The stronger *Stentor* has larger feeding flow. To compare the gains made by the weaker and the stronger *Stentor* by being in a pair, we define a new parameter the “the benefit of being together” for each *Stentor* as the difference between its feeding flow when in partnership with another *Stentor* and its feeding flow when alone, normalized by the latter, i.e.,

**Fig. 3:**
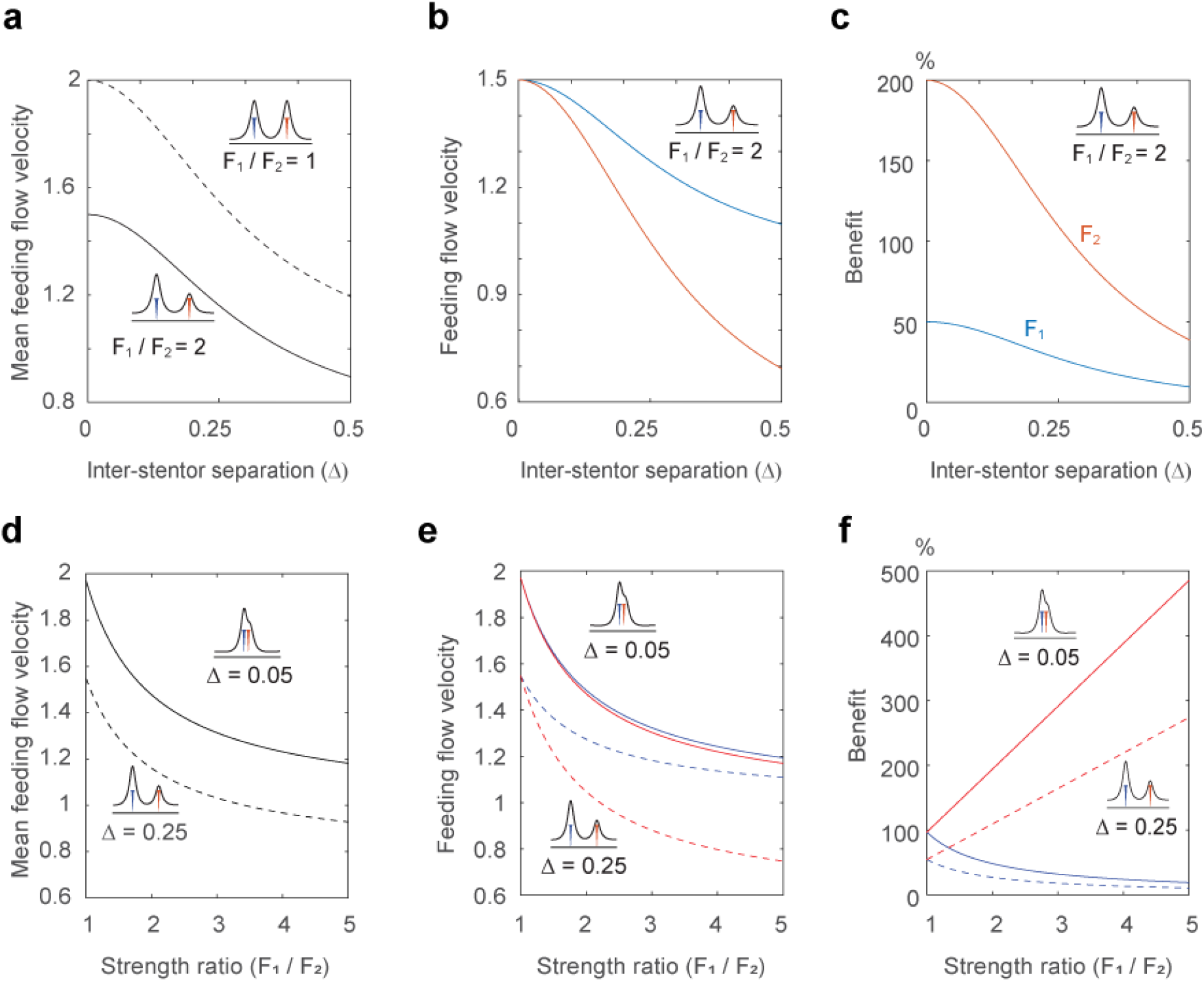
Mathematical modeling shows magnitude of combined feeding flow depends upon the inter-stentor separation Δ. **(a)** Average feeding flows for two *Stentor* pairs of different strength ratios *F*_1_/*F*_2_ = 1 and *F*_1_/*F*_2_ = 2. **(b)** Feeding flow of each *Stentor* in the *Stentor* pair with *F*_1_/*F*_2_ = 2. **(c)** The benefit of “being together” for each *Stentor* in the *Stentor* pair with *F*_1_/*F*_2_ = 2. Both *Stentor* benefit from their partnership, with increased benefit for the weaker *Stentor*. **(d)** Average feeding flows for two *Stentor* pairs at two different separation distance Δ = 0.05 (solid lines) and Δ = 0.25 (dashed lines). **(e)** Feeding flows of individual Stentors in each pair. **(f)** The benefit of “being together” for individual Stentors in each pair. The benefit to the stronger *Stentor* decreases as the strength of its partner decreases, irrespective of separation distance, while the benefit to the weaker *Stentor* increases multifold.

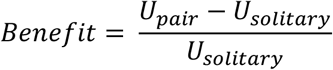

Our analysis clearly shows that both *Stentor* individuals benefit from their partnership. However, when the increased flows are compared relative to their feeding flow when alone, the weaker *Stentor* benefits more from the partnership (Fig. 3c). As discussed earlier, solitary *Stentor*s exhibit a wide dispersion in their flow velocities. We therefore conducted a similar analysis by varying the strength ratio of *Stentor*s (*F*_1_/*F*_2_) at two different inter-*Stentor* separation distances, Δ = 0.05 and Δ = 0.25 times body-length *H*. For all strength ratios, the pair with the smaller separation distance had higher average feeding flow (Fig. 3d). Moreover, the feeding flows of the stronger *Stentor* in a pair were always larger than the feeding flows of the weaker *Stentor* (Fig. 3e). However, the benefit to the stronger *Stentor* decreases as the strength of its partner decreases, irrespective of separation distance, while the benefit to the weaker *Stentor* increases as shown in (Fig. 3f). This pattern is consistent with our experimental results indicating that while both individuals benefit from being in close proximity, the advantage each receives is asymmetric with the weaker *Stentor* receiving a relatively higher benefit.

To date, benefits of colony formation have primarily been investigated using comparatively advanced organisms in which colony members are either embedded in the same matrix (e.g. *Volvox carteri*^3^) or physically attached to each other (e.g. *Zoothamnium dublicatum* and Choanoflagellate *Codosiga botrytis*^*26*^). Moreover, *Volvox* colonies consist of two differentiated cell types physically held together in an extracellular matrix. In contrast, *Stentor coeruleus* is a truly unicellular organism which exhibits a more basal colony forming behavior in which unit cells can reversibly aggregate to form dynamic multicellular-like colonies. Unlike *Volvox*, members of a *Stentor* colony are not physically connected to each other. Our results have demonstrated that proximity between individuals is sufficient for enhancement of feeding currents. Faster flows can potentially increase the rate of prey encounter and capture, and therefore provide colony members with a selective advantage over their solitary counterparts. This leads to the question: why would such an organism ever transition back to a solitary state when colonial organization has selective survival advantages? The answer may reflect the pattern among the genus *Stentor* to detach from their colonial organization and swim alone during periods of low prey abundance ^15^ or to avoid danger ^27^. While detachment from anchoring has been reported to reduce prey encounter rate by about 30 to 70% for some ciliates (*Pteridomonas danica* and *Paraphysomonas vestita*) ^28^, it enables rapid swimming and more effective maneuvering. These traits are valuable for cruising-mode foraging as well as in avoiding threats. In contrast, *Volvox* colonies can-not disassemble even under starvation conditions ^29^.

Our results agree with a recent theoretical study which predicted hydrodynamic cooperation between neighboring protists (Choanoflagellate) to increase fluid supply to the colony ^12^. For an isotropic colony with morphologically similar units, this study predicted that colony members would arrange themselves in a manner to maximize fluid flux. This would mean that once an optimal arrangement is achieved, their relative location in a colony would then remain unchanged to ensure continuation of maximal benefit. This is true in *Volvox* where the unit cells remain fixed on a uniform spheroid ^11^. Counter-intuitively, we observed that the separation between (oral openings of) members in a *Stentor* colony and in pairs oscillate over time, raising the speculation that this might be an inherent property of the organism. Further, differences between *Volvox* and *Stentor* colonies may be influenced by their physical organization. Unlike clonal members of a *Volvox* colony, *Stentor*s in a colony are not a homogeneous population and may include a variety of genotypes. Individuals across the same population show a wide dispersion in size as well as in the velocity of their feeding currents (Extended Data Fig. 2). These disparities between individuals can have major implications for their ability to get the most out of colony formation.

In this study, we found that paired *Stentor* individuals do not gain equally from the proximity i.e. the gains are asymmetric. If the pairings were permanent, one of the individuals would always be at a loss compared to the other, thus creating an imbalance. The dynamic nature of *Stentor* colonies (Fig. 4 a-c, Movie 1) might help address this imbalance by encouraging promiscuity between partners in a colony. We found that in a colony, while the nearest neighbor of each member frequently changes (Fig. 4 a,b), there is almost always a neighbor less than 0.25 mm away in a colony (Fig. 4c). Moreover, we found that about 86% of the time, each organism has a neighbor less than 0.025 mm away and 93% of the time a neighbor less than 0.05 mm away (Fig. 4c). This also explains why organisms in a pair do not always maintain the shortest possible separation to ensure maximal flow rates (Fig. 2). From an evolutionary standpoint, individuals should always be looking for the most favorable energetic payoff by associating with a neighboring individual that benefits them the most. One way to accomplish this is to keep switching between neighboring partners, particularly in a heterogenous environment when they are changing their location or height in a colony or going through contraction-extension cycles ^30,31^. Such asymmetric advantage could favor a dynamic pattern of colony members continuously changing their positions with respect to each other with the goal of maximizing their fluid flux under conditions of asymmetric advantages. We note that in a colony, even though an individual might appear to be moving away from one neighbor, it is actually moving closer to another neighbor (Fig. 4a). Although *S. coeruleus* appears to be a less-advanced organism than *Volvox*, the ability of colony members to dynamically alter their morphology and location endows it with very unique benefits. Our experiments demonstrate that the reversible and dynamic nature of *S. coeruleus* colonies might represent an initial pathway from isolated individuals to coordinated colonial activities.

**Fig. 4:**
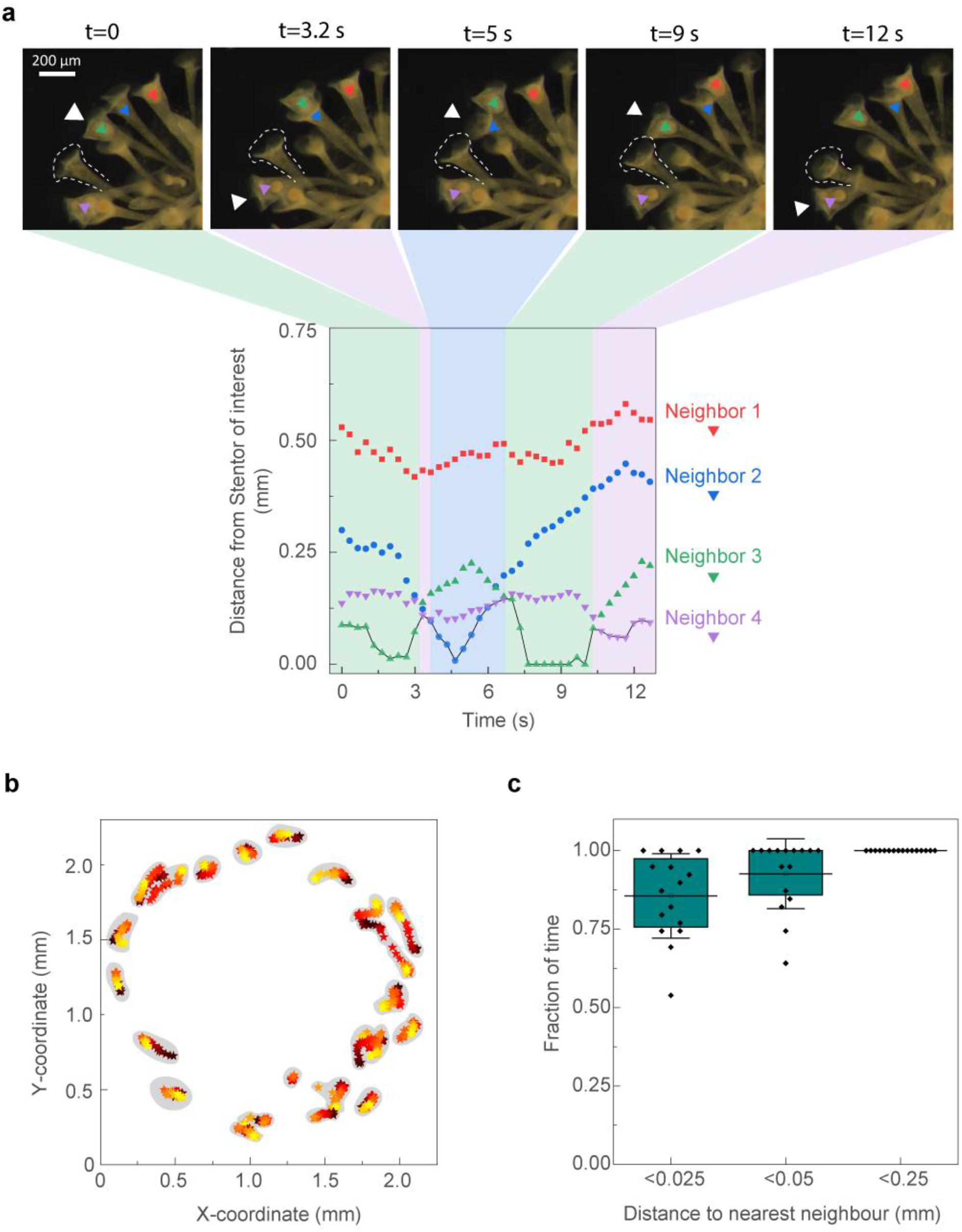
Dynamic relocation of individuals in a colony ensures constant presence of proximal neighbors. **(a)** Top: dark-field images showing movement of 4 neighboring Stentors from a colony of wild *Stentor muelleri* (neighbors are indicated by red, blue, green and magenta arrowheads) with respect to a specific individual (denoted by dotted outline) in a colony. The 5 Stentors are in the same focal plane. The white arrowhead indicates the closest neighbor to the individual of interest. Below: The distance of each of the neighbors (colored arrowheads above) from the *Stentor* of interest (dotted outline above) over time. The closest neighbor to the individual of interest changes from Neighbor 3 to, Neighbor 4 to Neighbor 3 to Neighbor 4 over the course of time (denoted by the black thick line). **(b)** The time-dependent location of the center of the oral apparatus (head) of individuals in a colony recorded in a single focal plane over 13 seconds (see Movie 1, Fig. 1e). Each cluster represents the time-dependent location of an individual. Symbols in a single cluster are colored to represent time (black→ red → yellow from t = 0 to 13 seconds). **(c)** Fraction of total time an individual in the above colony has at least one neighbor <0.025 mm, 0.05 mm or 0.25 mm away. Horizontal line is the mean, error bars represent standard deviation (n=16) and the box represents 25/75th percentiles.

## Methods

### *S. coeruleus* and *S. muelleri* culturing

*Stentor coeruleus* cells were obtained commercially (Carolina Biological Supply, Burlington, NC) and were subsequently cultured as described earlier ^32^. Briefly, cells were grown in the dark at 20°C in Modified *Stentor* Medium (MSM) (0.75 mM Na_2_CO_3_, 0.15 mM KHCO_3_, 0.15 mM NaNO_3_, 0.15 mM KH_2_PO_4_, 0.15 mM MgSO_4_, 0.5 mM CaCl_2_, 1.47 mM NaCl) modified from recipes by Tartar ^15^ and De Terra ^33^. This medium was supplemented with living prey, *Chlamydomonas reinhardtii* which were washed in MSM before being added to the *S. coeruleus* cultures for feeding. A few grains of boiled wheat seeds were also added to the *S. coeruleus* cultures to promote additional microbial growth. *C. reinhardtii* cells were added to the *S. coeruleus* cultures two or three times per week.

*S. muelleri* were taken from Shivericks Pond in Falmouth, MA, USA. About 2 liters of pond water along with organic matter like leaves and twigs were collected from a shaded part of the pond. The sample was brought back to the lab, mixed vigorously by shaking the bottles, and then allowed to settle for about 10 minutes. The water containing the freely swimming organisms was then separated from the organic matter and poured into a borosilicate glass flask. *S. muelleri* were identified under a stereo microscope and isolated using a P1000 pipette into a separate flask. Filtered pond water was added to the culture and S. *muelleri* were used in the experiments in the next 24 hours.

### Microscopy, data acquisition and analysis

Stentors suspended in water mixed with whole milk (diluted 500x) were incubated in a 5 mm high PDMS chamber until they anchored themselves on the glass coverslip attached to one end of the PDMS. Once anchored on the coverslip, the Stentors were imaged in dark field using a 4x objective on an inverted microscope (Nikon Eclipse TE2000-U). For flow visualization, dark field time-lapsed images of the anchored Stentors with milk particles in their surrounding liquid were acquired at 500 frames per second using a high-speed color camera (Fastcam 1024 SA3; Photron), such that 1 pixel on the camera corresponded to 4.2 μm.

Median images were calculated for each video using Fiji and were subtracted from individual video frames to eliminate particles stuck on the surface. These processed images were Z-projected using Fiij to generate images of flow streamlines (Fig. 1c,f). To determine velocity and vorticity fields, these median-subtracted videos were analyzed with a cross-correlation algorithm using DaVis 8.0 (LaVision, Germany). Image pairs were analyzed with shifting overlapping interrogation windows of a decreasing size of 32 × 32 pixels to 16 × 16 pixels. Once the velocity field was measured, the velocity of the feeding current was determined by calculating the average velocity across a plane the same size of the oral opening at a distance of 0.25 mm from the oral opening (Extended Data Fig. 1).

### Mathematical modeling of feeding flows

Following common practice in modeling sessile micro-organisms, we consider the details of the cilia length, beating frequency, and waveform of the *Stentor’s* ciliated ring to be subsumed into a single averaged force ***f*** that the *Stentor* exerts on the fluid. Specifically, we model each *Stentor* as a regularized Stokeslet force ***f***= −*F****e***_3_ = (0,0, −*F*) located at ***x***_o_ = *H****e***_3_ = (0,0, *H*), that is, at a height *H* above a solid flat wall at z = 0. Here, without loss of generality, we considered Cartesian coordinate (*x, y*, z) located at the base (attachment point) of the *Stentor*, with ***e***_3_ the unit vector in z-direction. We set the regularization parameter *a* to be 0.1*H*, which roughly corresponds to the radius of the ciliated ring.

The fluid flow generated by the *Stentor* in the model is governed by the incompressible Stokes equations with zero boundary conditions at the solid wall z = 0 and at infinity, namely,

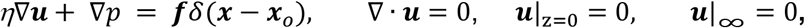

where ***u*** is the fluid velocity, *p* the pressure field, *η* the viscosity of the fluid, and δ the three-dimensional Dirac-Delta function.

To solve for the fluid flow field ***u***, we use the regularized Stokeslet method proposed by Cortez^24^. The no-slip wall is achieved by adding an appropriate image system to the Stokeslet. The image system was originally derived by Blake ^22^ and was recently reformulated by Gimbutas et al.^34^. The regularized version of the image system was studied by Ainley et al. ^35^, based on Blake’s formulation. The image system consists of the regularized counter-parts of a Stokeslet, a potential dipole, a Stokes doublet, and a rotlet. In general, the fluid velocity at the point ***x***_*e*_ induced by a force ***f***located at ***x***_o_ = (*x*_o_, *y*_o_, z_o_) and its images at ***x***_o,*im*_ = (*x*_o_, *y*_o_, −z_o_) is given by

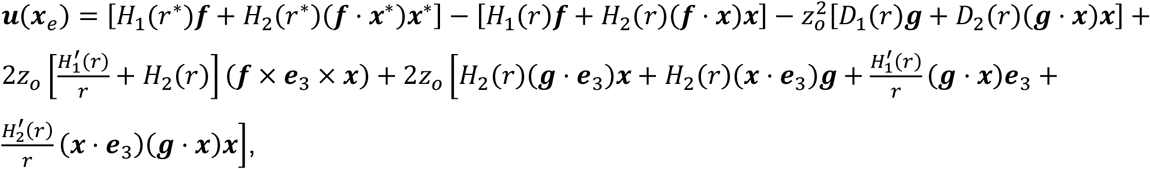

where ***x***^∗^ = ***x***_*e*_ − ***x***_o_, ***x*** = ***x***_*e*_ − ***x***_o,*im*_, *r*^∗^ = |***x***^∗^|, *r* = |***x***|, ***g*** = 2(***f***⋅ ***e***_3_)***e***_3_ − ***f*** is the dipole strength, and 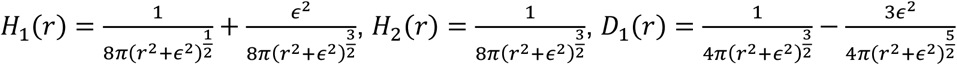, and 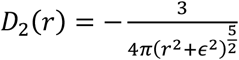.

Note there is a typo in the original paper that we have now corrected in the above expression. We refer interested readers to the aforementioned papers for the derivation of these expressions. A typical flow field of the model *Stentor* for the parameters listed in Extended Data Table 1 is shown in Extended Data Fig. 4. Here, the Stokeslet force scaled as *F* ∼ *ηUH*.

We next considered a pair of Stokeslet, representing a pair of Stentors located side-by-side as detailed in the main document. To study the asymmetry between the *Stentor* pair, we kept the strength of the left *Stentor* (*F*_1_) to be constant and decrease the strength of the right *Stentor* (*F*_2_) so that 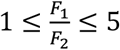

To quantify the feeding flow velocity *U*, we measured the average downward velocity component over the area of the ciliated ring π*a*^2^ at a distance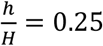 from the ciliated ring of each *Stentor*. Specifically, *U*_*s*o*litary*_ is measured when there is a single *Stentor* in the system and *U*_*pair*_ is measured for each *Stentor* when there are two Stentors placed side-by-side in the system.

## Data and code availability

Data supporting the findings of this manuscript and code are available from the corresponding authors upon reasonable request.

## Acknowledgements

This work was supported by NIH NIGMS grant R35GM130327 to WM, Whitman Early Career Award from MBL, NIH NIGMS R35GM143050 and startup funds from Emory University to SS.

## Author contributions

S.S. and J.H.C. designed the experiments, S.S. conducted the experiments, S.S., S.P.C and W.M. analyzed the data, H.G. and E.K. designed the mathematical model, S.S., J.H.C, H.G. and E.K. wrote the manuscript.

## Note

This is a preprint and is not peer reviewed.

## Competing interests

We declare no conflicts of interest.

## Correspondence

and requests for materials should be addressed to S.S., J.H.C. and E.K.

## Notes

### Competing Interest Statement

The authors have declared no competing interest.

