## Supplementary figures and table for "Cooperative hydrodynamics accompany multicellular-like colonial organization in the unicellular ciliate *Stentor*"

**This PDF file includes:**

Extended Data Figs. 1 to 4

Extended Data Table 1

### Extended Data Figures

**Extended Data Fig. 1: Schematic representation of the methodology for measurement of feeding currents of *S. coeruleus*.**

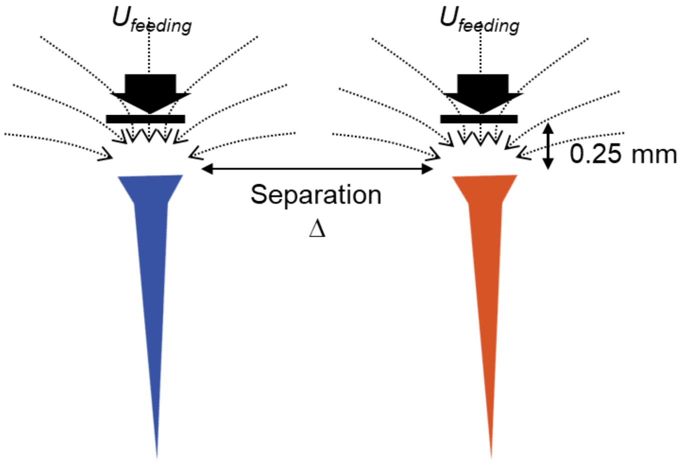


Since only the orthogonal portion of the flowfields in the region between the two vortices reaches the oral opening and can be productively filtered for prey, only the velocity component normal to the oral opening is taken as a measure of the feeding current. Due to mixing between tracer and cilia movements, feeding current velocities were measured 0.25 mm from the oral opening. Feeding current for each individual in a pair or alone was determined by calculating the average velocity across a plane the size of the oral opening of each individual.

**Extended Data Fig. 2: Velocities of feeding currents of solitary *Stentor* individuals show a broad distribution**

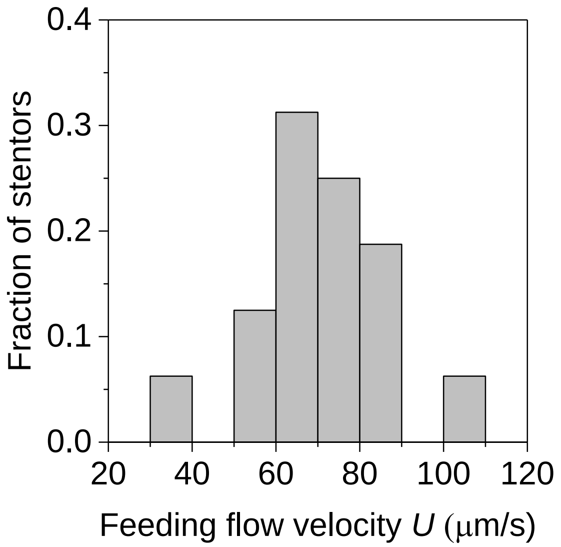


Distribution of velocities of the feeding current generated by solitary *Stentor* individuals (n=17) measured at 0.25 mm from the ciliary band.

**Extended Data Fig. 3: Velocity of the ciliary wave remains unchanged as the proximity between a pair changes.**

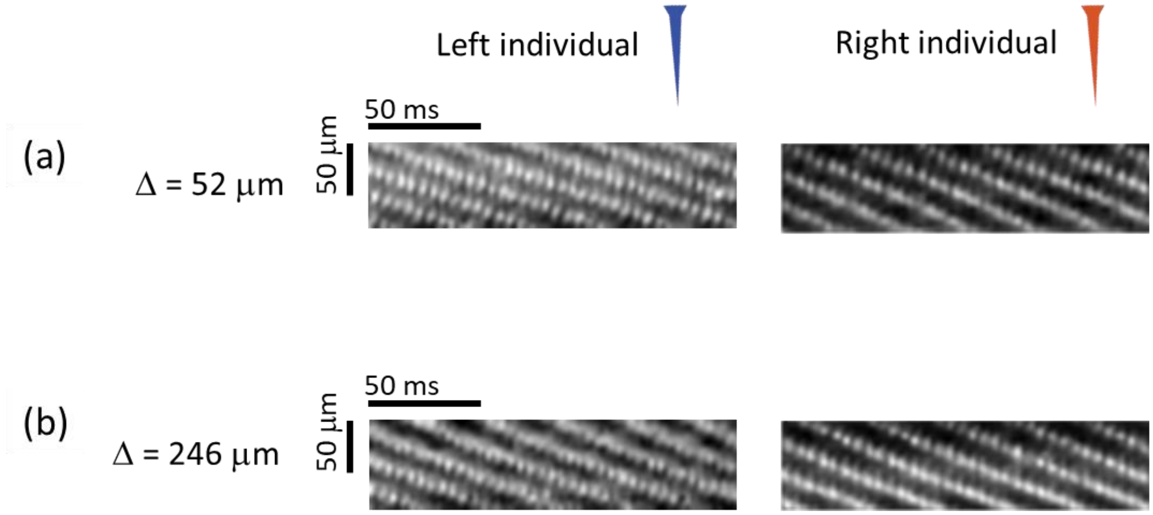


Kymographs showing the frequency of the metachronal waves for the two individuals in the pair shown in Fig. 3a,b. The frequency of the metachronal wave for the left and the right *S. coeruleus* in the pair remains unchanged as the distance between the two individuals changes from **(a)** Δ = 52 μm (left: 16.7 Hz, right: 15.4) to **(b)** Δ = 246 μm (left : 16.6 Hz, right: 15.2 Hz).

**Extended Data Fig. 4: Streamlines generated by the model *Stentor*.**

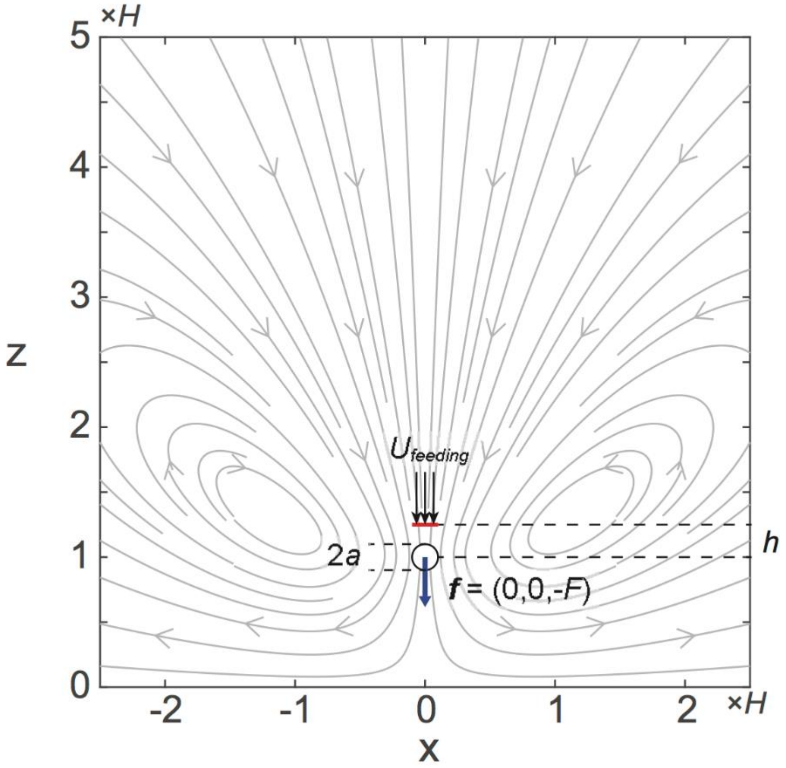


The *Stentor* is modeled by a regularized force pointing towards a no-slip wall. The force is located at a distance $H$ from the wall and the regularization parameter is set to be $H/10$. The feeding velocity $U$ is the flux going through a circular cross-section of radius $a = H/10$ at a height $h=H/4$ from the force, averaged over the circular area.

**Extended Data Table 1: Characteristic scales of the Stokeslet model**

| Parameter | Symbol | Dimensional value |
|---|---|---|
| *Stentor* height | $H$ | 1 mm |
| Fluid viscosity | $\eta$ | $10^{-3}$ Pa s |
| Flow velocity | $U$ | 100 µm $s^{-1}$ |
